# Epigallocatechin-3-Gallate Improves Facial Dysmorphology Associated with Down Syndrome

**DOI:** 10.1101/276493

**Authors:** John M. Starbuck, Sergi Llambrich, Ruben González, Julia Albaigès, Anna Sarlé, Jens Wouters, Alejandro González, Xavier Sevillano, James Sharpe, Rafael de La Torre, Mara Dierssen, Greetje Vande Velde, Neus Martínez-Abadías

## Abstract

In Down syndrome (DS), the overall genetic imbalance caused by trisomy of chromosome 21 leads to a complex pleiotropic phenotype that involves a recognizable set of facial traits. Several studies have shown the potential of epigallocatechin-3-gallate (EGCG), a green tea flavanol, as a therapeutic tool for alleviating different developmental alterations associated with DS, such as cognitive impairment, skull dysmorphologies, and skeletal deficiencies. Here we provide for the first time experimental and clinical evidence of the potential benefits of EGCG treatment to facial morphology. Our results showed that mouse models treated with low dose of EGCG during pre- and postnatal development improved facial dysmorphology. However, the same treatment at high dose produced disparate facial morphology changes with an extremely wide and abnormal range of variation. Our observational study in humans revealed that EGCG treatment since early in development is associated with intermediate facial phenotypes and significant facial improvement scores. Overall, our findings suggest a potential beneficial effect of ECGC on facial development, which requires further research to pinpoint the optimal dosages of EGCG that reliably improve DS phenotypes. Current evidence warns against the non-prescribed intake of this supplement as a health-promoting measure.

## Introduction

The face, despite being a variable phenotype with high levels of plasticity and remodeling throughout the life of an individual, requires the fine orchestration of multiple morphogenetic events during early development and subsequent growth^1^. Changes to this long-term program can impair growth and bone remodeling and result in facial shapes that fall beyond the typical range of facial morphological variation^2^. Facial dysmorphologies are often associated with congenital and neurodevelopmental disorders. Since brain and face are intimately integrated^3^, genetic and/or environmental insults in early embryogenesis can alter the development of both structures. A major genetic alteration results from aneuploidies, which involve the presence of an abnormal number of whole or part of a chromosome^4^. Trisomy 21 (Ts21), referred to as Down syndrome (DS) (OMIM 190685), is a complex pleiotropic genetic disorder that is the most common genetic cause of intellectual disability and is associated with a characteristic facies. The specific facial features of DS (i.e., midfacial flatness, oblique palpebral fissures, mandibular hypoplasia, and facial asymmetry), may cause facial stigmata and compromise quality of life^5^.

Besides altering neural^6^ and facial development^7^, the overall genetic imbalance resulting from the overexpression of hundreds of chromosome 21 genes perturbs the morphogenesis and growth of many other body systems and organs, such as immune^8^ and gastrointestinal^9^ systems, as well as the heart^10^ and the skeleton^11^. The complex and widespread phenotypic outcomes of DS are thus the result of combined effects of genes altering specific tissues^12^, as well as the integrated effect of genes altering common signaling pathways simultaneously in different tissues during development. This genetic complexity, combined with large degrees of developmental and phenotypic variability in individuals with DS, represents a serious challenge to any attempt to alleviate the alterations caused by Ts21.

Nevertheless, recent studies have shown that epigallocatechin-3-gallate (EGCG), a green tea flavanol, is a promising therapeutic tool for individuals with DS^13–17^. Clinical studies indicate that EGCG doses equivalent to a daily intake of eight cups of green tea show efficacy in improving adaptive functionality and cognitive deficits associated with DS^13,16^. Similar effects have been found in a study using DS mouse models^13^. EGCG is a well-known kinase inhibitor^18^, but the optimal dosage, onset and duration of the EGCG treatment in DS remain to be elucidated^19^.

The potential of EGCG as an effective treatment for DS relies, at least in part, on its ability to inhibit the gain of function of the dual specificity tyrosine-phosphorylation-regulated kinase 1A (*DYRK1A)*^20^, which is one of several chromosome 21 genes triplicated by Ts21 and at dosage imbalance in individuals with DS that map to the so-called Down syndrome critical region (DSCR)^21^. Traditionally, the activity of *DYRK1A* has been associated with neural development^22^, but in recent years new roles for *DYRK1A* have been identified for development of the face^23^, the skeleton^14,15^, and the heart^24^. Interestingly, recent studies in mice have reported that EGCG may alleviate some skull^17^ and skeletal alterations^14,15^ in DS mouse models.

Given the potential of EGCG as an effective treatment to minimize many of the alterations present in DS^13,15–17^, we here focused on assessing the effect of EGCG on the development of the face. We started examining the effect of a standardized EGCG treatment on facial skeletal development using the well-established *Ts65Dn* DS mouse model. We also performed an observational study to investigate for the first time whether EGCG treatment also affects facial dysmorphology in children with DS.

### Experimental effect of EGCG treatment on facial shape in DS mouse model

We explored the potential of EGCG prenatal treatment to prevent the facial malformations associated with DS, assessing facial morphology in trisomic (TS) and euploid (wild-type or WT) mice after randomly treating half of the litters either with a high (30 mg/kg/day) or a low (9 mg/kg/day) EGCG dose from embryonic (E) day 9 to post-natal (P) day 29 (see Table SI_1 for sample composition and Materials and Methods for more details on mouse models and treatment conditions). We compared mouse facial shape using the 3D coordinates of 12 facial landmarks registered on micro-computed tomographic (μCT) images of the craniofacial skeleton of the mice (Fig. SI_1 and Table SI_2).

Results of a Procrustes-landmark^25^ based shape analysis showed that, despite slight morphospace overlap, untreated WT and TS mice showed different facial shapes (Fig. 1). However, the effect of the EGCG treatment varied greatly depending on the dose. Both WT and TS mice treated with the high EGCG dose presented large ranges of morphological variation, spanning through the whole morphospace (Fig. 1). Regardless of their genotype, mice presented disparate facial phenotypes: some mice, both WT and TS, fell within the typical range of variation of WT untreated mice, whereas others fell on the opposite extreme of variation, associated with unique morphology and severe facial phenotypes (Fig. 1). These results indicated that, at high dosages, the effect of EGCG on facial shape is highly variable and potentially detrimental in some cases.

**Figure 1.**
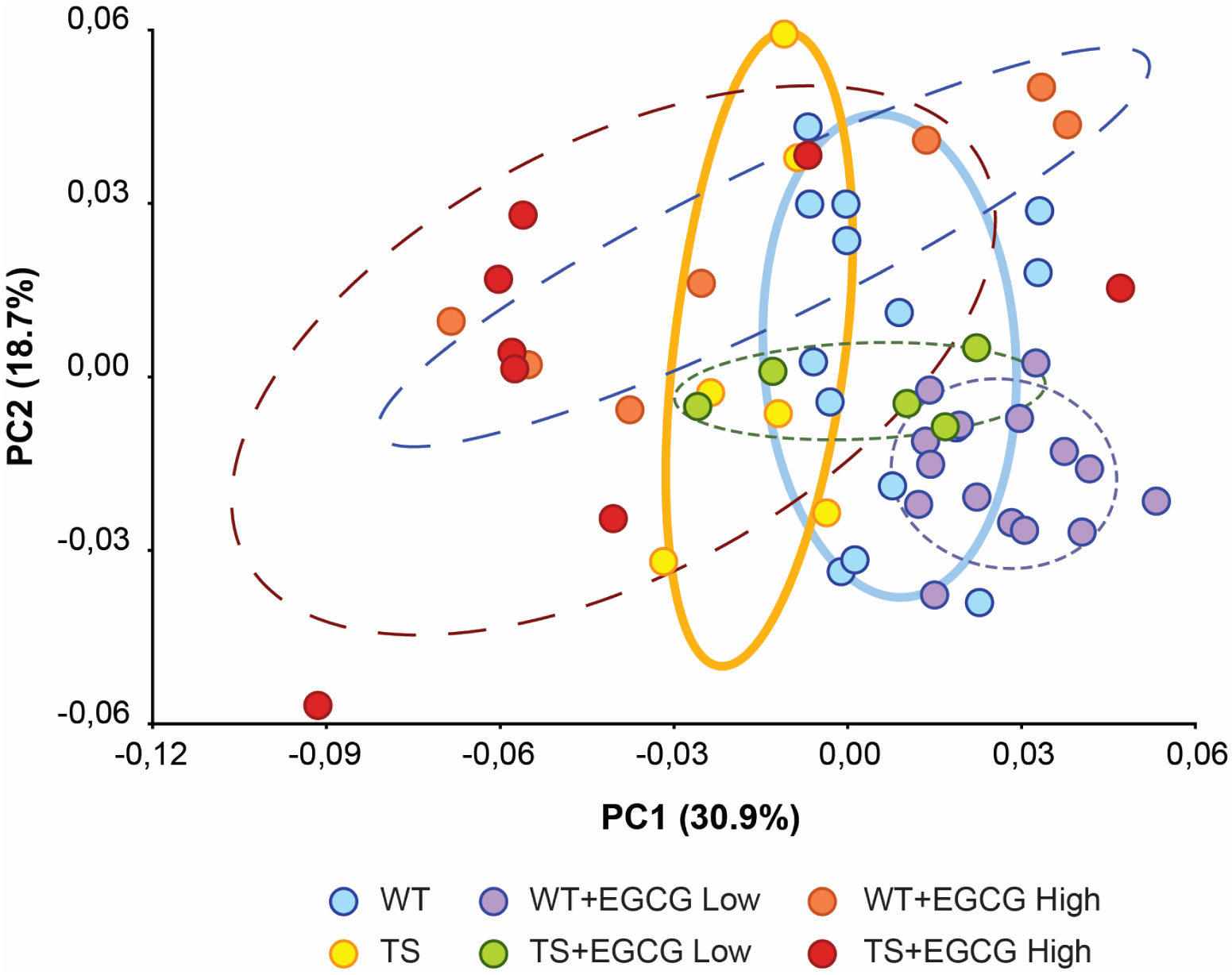
Principal Component Analyses based on the Procrustes-based landmark analysis of the global facial shape in DS mouse models treated at high and low EGCG dose. Scatterplot of PC1 and PC2 axes with the corresponding percentage of total morphological variation explained is displayed. Ellipses represent the 70% confidence interval within each group of mice.

On the contrary, mice treated with the low EGCG dose showed moderate ranges of variation and both WT and TS treated mice presented WT-like facial shapes, different from TS facial shapes (Fig. 1). WT mice treated at low EGCG doses did not show any effect of the treatment on facial shape, whereas 80% of the TS mice treated with low EGCG doses fell within the range of variation of untreated WT mice (Fig. 1). This strongly supports a predictable normalization of global facial shape upon low dose EGCG treatment.

To provide further evidence about the effects of EGCG on murine facial morphology at a more localized level, we developed a novel approach to estimate facial improvement scores (FISs) and assess if the EGCG treatment significantly changes the number of facial traits that differentiate the euploid and the trisomic facial phenotypes for each of the EGCG dosages. This approach is based on the results of Euclidean Distance Matrix Analyses (EDMA)^26^, a technique that calculates all unique linear distances between the facial landmarks for each individual and performs pairwise shape comparisons between groups. To estimate FIS scores, we first ran an EDMA to detect how many linear distances were significantly different between WT and TS untreated mice. Then, we ran another EDMA between WT mice and TS mice treated with EGCG, and by comparing the results from both EDMAs we estimated the corresponding FIS (see formula in Materials and Methods). A FIS score close to 0 represents no effect of the EGCG treatment, as the number of significant facial differences between WT and TS mice remains equal with or without EGCG treatment. A positive FIS score represents a normalization of the facial phenotype, as it is achieved by a reduction of the facial shape differences between WT and TS mice after treatment. A negative FIS score indicates a detrimental effect of the treatment, in which the EGCG treatment exacerbates the severity of the DS facial traits, increasing the number of facial shape differences between WT and TS mice. Finally, to confirm that the patterns revealed by the EDMA analyses were not due to random chance alone, we tested the statistical significance of the results performing simulation tests. We performed a subsampling approach in which we generated random pseudo-subsamples with increasing number of DS treated cases, and for each simulation we recomputed all the EDMA analyses and estimated the associated FIS scores (see Materials & Methods for further details).

The histograms in Fig. 2 compare the distribution of randomly generated FIS scores with the FIS obtained using the experimental data. Although the tests did not reach statistical significance because of large degrees of variation, the results of the simulation analyses tended to confirm our previous results: a negative effect of the high EGCG dose, but a positive effect of the low dose. The high dose EGCG treatment was associated with a negative FIS, as the number of facial differences between WT and TS mice increased when more TS treated mice were included in the simulated groups, and thus suggesting a detrimental effect of the high dose EGCG treatment (Fig. 2). On the contrary, the low dose EGCG treatment was associated with a positive FIS (Fig. 2), as the number of facial differences decreased with increasing number of TS treated mice, suggesting a normalization of the facial phenotype at low EGCG doses.

**Figure 2.**
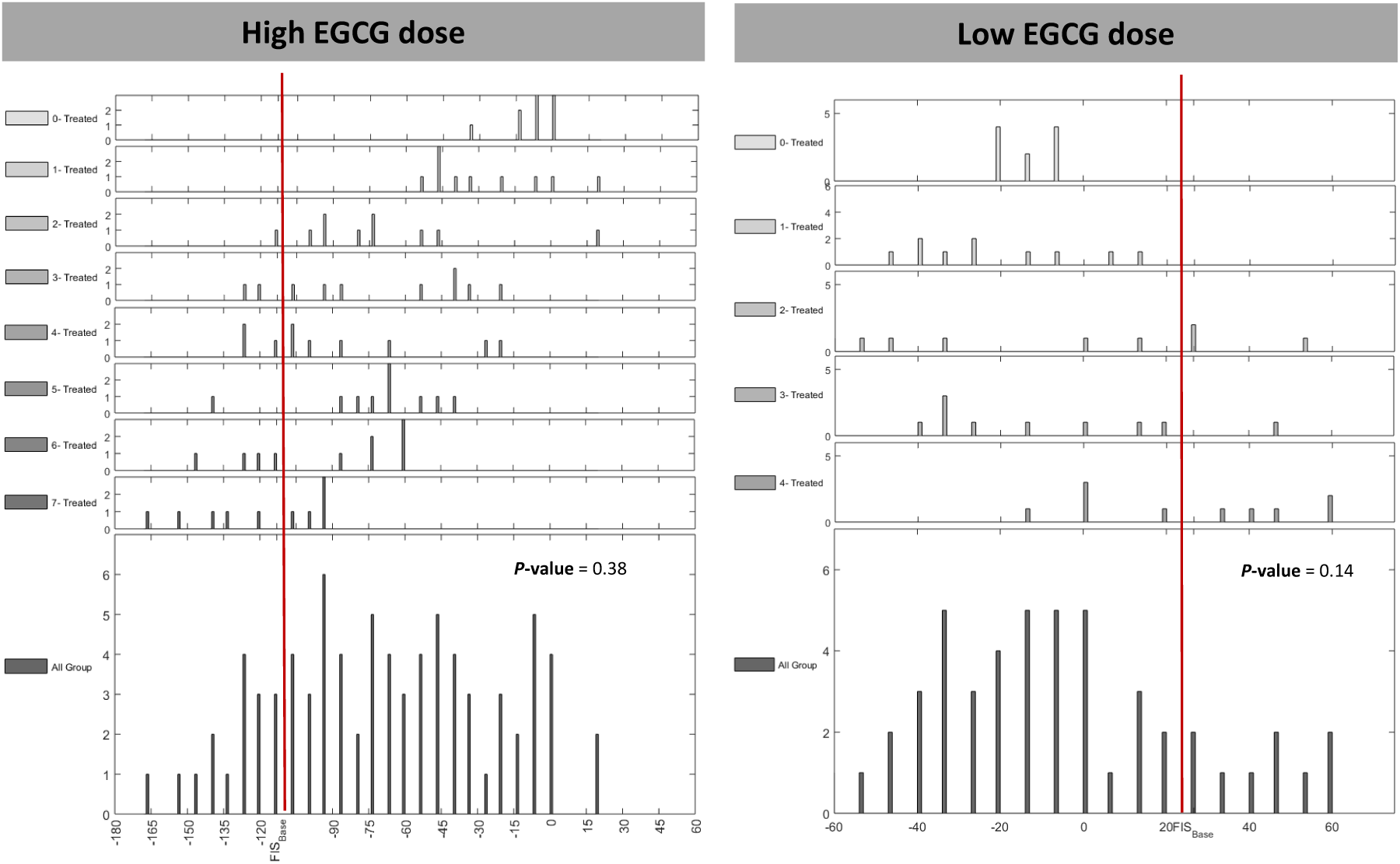
Simulation tests based on facial improvement scores (FIS) in DS mouse models that received an experimental prenatal EGCG treatment. FIS is a summary value that was computed by contrasting the number of significant differences in facial traits between 1) wildtype mice and TS mice not treated with EGCG and 2) wildtype mice and TS mice treated with EGCG. Histograms representing the simulation results for each random group are provided separately as well as aggregately. Each group contains an increasing number of EGCG treated mice. The red line shows the FIS score obtained with the observed EGCG treated mice. *P-value*s are provided for each group.

### Effect of EGCG treatment on facial shape of children with DS

The findings based on the experimental DS mouse models led us to explore whether EGCC treatment can also modulate facial development in humans. To this aim, and given that some parents are administering over-the-counter green tea extracts to their children because these are commercially available as dietary supplements, we were able to perform the first observational study ever to investigate whether EGCG treatment improves facial dysmorphology in children with DS. Our investigation is a cross-sectional study, on a cohort of individuals with DS that started green tea consumption on the initiative of their parents encouraged by the pro-cognitive effects of EGCG in DS. The participants in our study did not follow a systematic EGCG usage pattern, and thus had not homogeneous or standardized dosage, onset, and duration of treatment. Yet, assessing whether there is an effect of EGCG treatment on DS facial development may provide valuable evidence to determine whether EGCG treatment can alleviate facial dysmorphologies in DS.

To assess the effect of EGCG supplementation on facial development of children with DS, we first acquired 3D facial images of children with DS, children with DS that have received EGCG treatment, mosaic cases with quantified percentages of trisomic cells (50-80%), and euploid individuals (see Table S1_3 for sample composition and Table SI_4 for cohort and EGCG treatment details). Individuals were categorized into three age groups (0-3 yrs., 4-12 yrs., 13-18 yrs.) to determine whether EGCG treatment could have differing effects depending of the stage of development and onset of the treatment. Hence, facial shape was evaluated within each age group using multivariate geometric morphometric methods. We first assessed the overall face shape differences between euploid children and children with DS, with particular focus on children with DS that had received EGCG supplementation. Our results, based on Procrustes analyses of a configuration of 21 landmarks located throughout the face (Fig. SI_1), showed almost complete separation between euploid children and those with Ts21 in all three age groups (Fig. 3), with slight to moderate overlap depending on the specific age range. Overall, facial phenotypes associated with DS included flat and wider faces, oblique palpebral fissures, flat nasal bridges, wider noses, and thicker lips. Facial phenotypes associated with the euploid condition presented slender and slightly convex faces, with horizontal palpebral fissures, narrow noses and thin lips. The position in the morphospace of the DS cases treated with EGCG varied greatly in the different age groups. Depending on the age of initiation of the treatment, DS cases treated with EGCG showed facial phenotypes that fell within the ranges of variation of either the DS or the euploid groups (Fig. 3).

**Figure 3.**
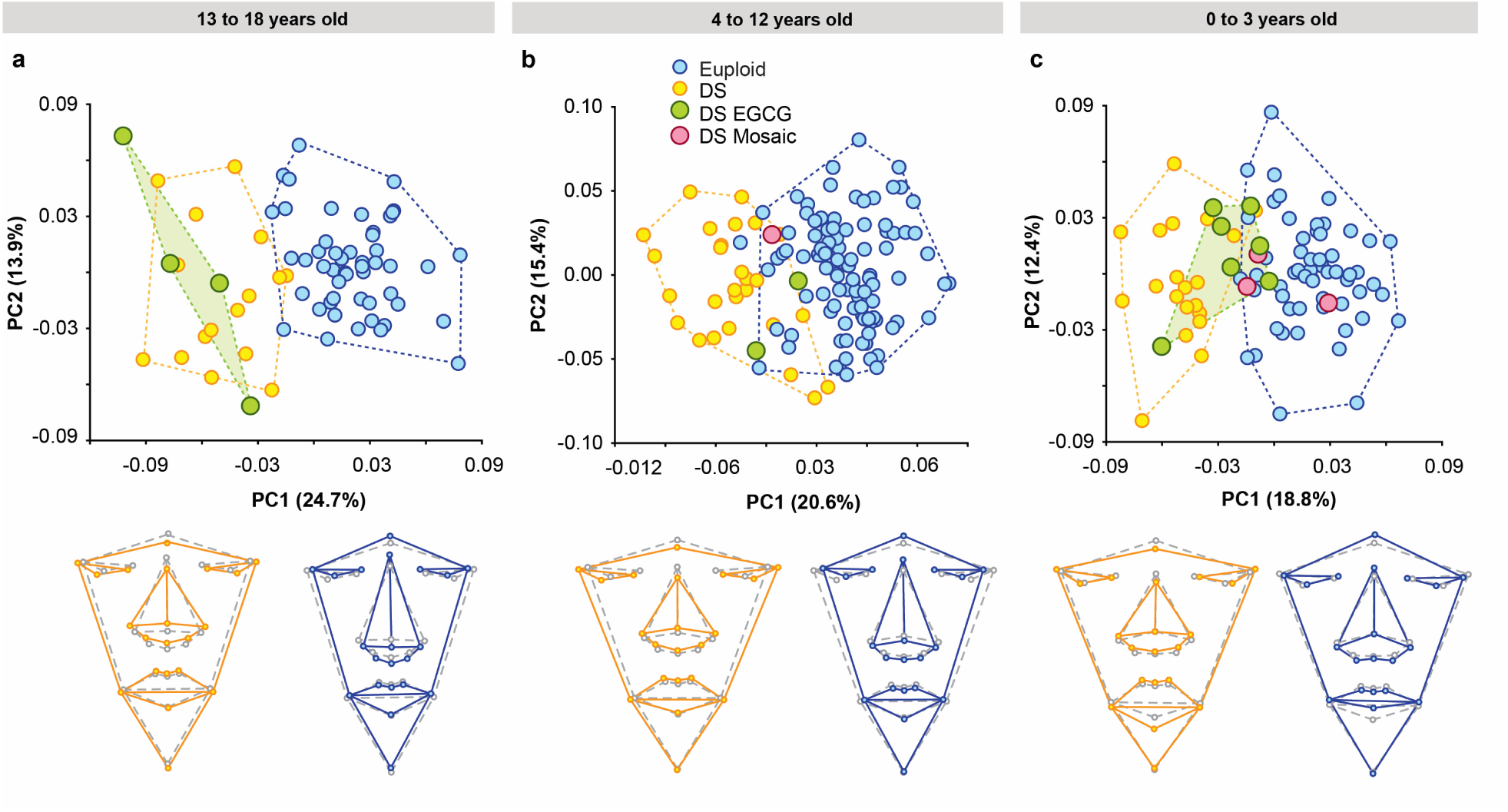
Principal Component Analyses based on the Procrustes-based landmark analysis of the global facial shape at each age group. a) Children from 13 to 18 years old, b) Children from 4 to 12 years old, c) Children from 0 to 3 years old. Scatterplots of PC1 and PC2 axes with the corresponding percentage of total morphological variation explained are displayed for each analysis, along with anterior facial morphings associated with the extreme negative and positive values of PC1. Orange solid line represents facial phenotype associated with DS, whereas blue solid line represents facial phenotype associated with euploid condition. Both shapes are compared to average facial shape, represented by dashed grey line. Convex hulls represent the ranges of variation within each group of children.

In the age group including teenagers from 13 to 18 years old, the participants that had taken the EGCG treatment completely overlapped with other individuals with DS that never received a treatment. Thus, the facial shapes of teenagers with DS were similar regardless of the EGCG treatment, suggesting that EGCG did not have any effect on facial shape when the treatment started late in adolescence (Fig. 3a). In the age group from 4 to 12 years old, the two DS treated cases fell in an intermediate morphospace position, one within the euploid range of variation and the other within the overlapping area between the euploid and the DS range of variation. This result showed a tendency of the children with DS treated with EGCG to display milder facial phenotypes. The child receiving the lower dose fell closer to the euploids, but no conclusive evidence can be drawn from these results due to the small sample size and the substantial differences in the dosage/length of EGCG treatment (Fig. 3b).

Finally, in the age group of youngest children (from 0 to 3 years old), we observed the most prominent effect of EGCG treatment: most children with DS who had taken EGCG since early in development presented a moderate facial phenotype (Fig. 3c). Six out of seven EGCG treated cases fell in an intermediate morphospace position between the euploid and trisomic samples. Three fell in the range of variation of children with DS showing mild facial phenotypes closer to euploid morphospace, and three fell within the range of variation of the euploid subsample (Fig. 1c). Only one child with DS treated with EGCG did not follow this tendency and was positioned within the range of variation defined by untreated DS cases (Fig. 3c), but this difference could not be significantly associated with any of the conditions of the treatment.

The position of three mosaic DS individuals within the range of variation of euploid children correlated with the degree of mosaicism. The mosaic case with the lowest reported percentage of trisomic cells (50%) fell in the middle of the range of variation of euploid children, whereas the other two mosaic cases with higher percentage of trisomic cells (75 and 80%) showed a more typical DS phenotype, but still overlapped with euploid children and DS cases with less marked DS facial traits (Fig. 3c). Our results thus provide evidence that reduced degree of genetic imbalance, resulting either from mosaic trisomy or EGCG treatment, is associated with milder facial phenotypes and reduced dysmorphology.

Next, we used EDMA to detect statistically significant local facial shape differences and to pinpoint exactly where EGCG may influence facial shape. We estimated the baseline comparison of DS and euploid samples, and then assessed whether the result for each linear distance was significantly different when the comparison only included children with DS that had been treated with EGCG. A potential beneficial effect of EGCG would be revealed by a decrease in the number of significantly different facial measurements between euploid and trisomic children treated with EGCG relative to the baseline comparison of the euploid children and untreated DS cases.

When assessing facial differences in the 13-18 year age group, our results indicated only a small reduction in the number of significant linear distances. A total of 61% of linear measures significantly differed between the baseline comparison of DS and euploid samples (Fig. 4a), while 53% were still significantly different between euploid and treated samples (Fig. 4b). When comparing untreated children with DS and those treated with EGCG, only a minimal number of significant differences were found (5%), indicating overall similarity between untreated and treated children with DS (Fig. 4c, d).

**Figure 4.**
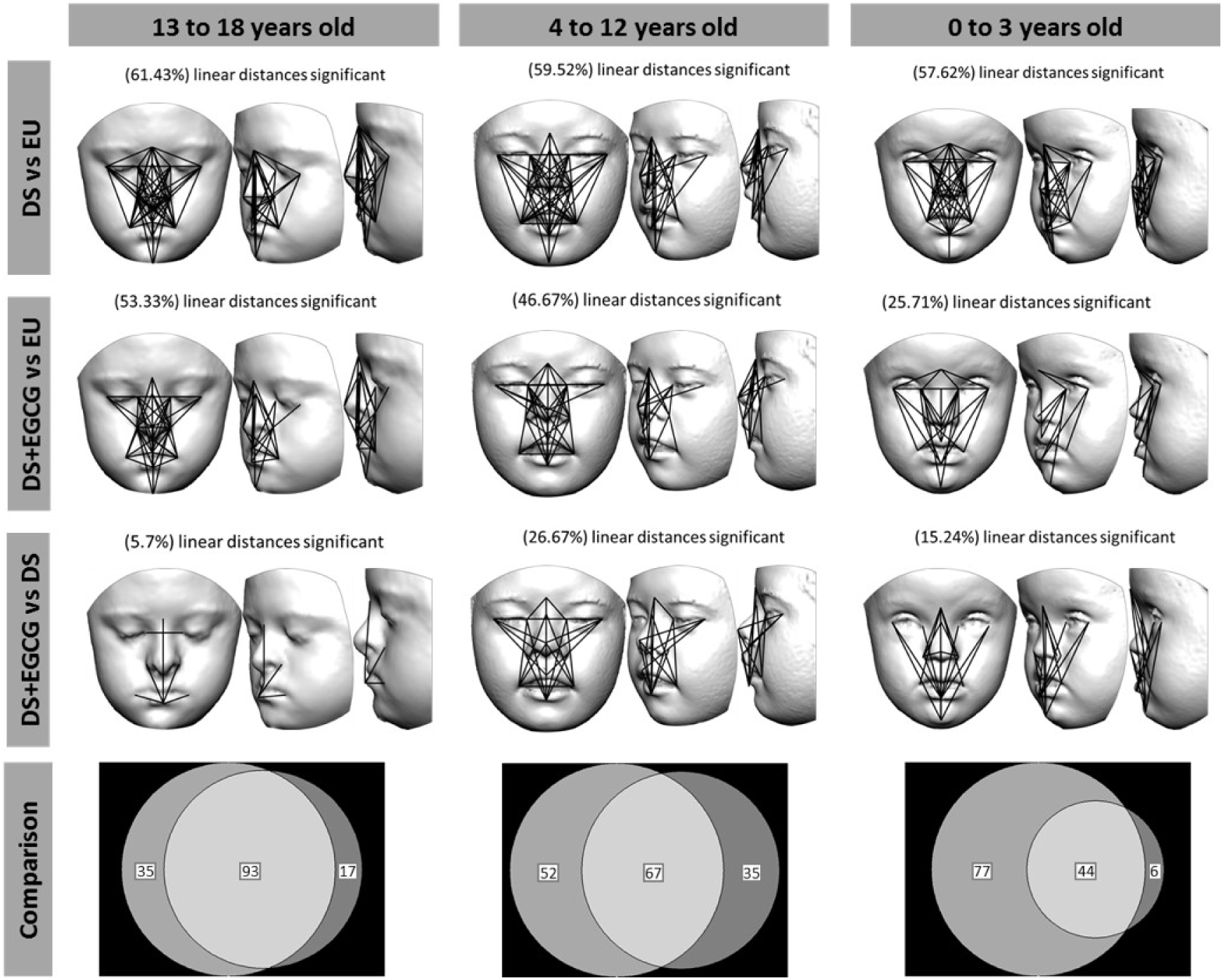
Localized facial shape comparisons based on EDMA analyses at each age group. Black solid lines represent linear facial measurements that are significantly different in the two groups being compared. First row: children with Down syndrome (DS) *versus* euploid children (EU); second row: children with Down syndrome treated with EGCG (DS+EGCG) *versus* euploid children (EU), third row: EGCG treated (DS+EGCG) *versus* non treated children with Down syndrome (DS), last row: Venn diagrams representing number of significant differences that are common or unique in each comparison for each age group. Specifically, the circles in the Venn diagrams indicate the number of unique significant differences DS *vs* EU (left), the number of common significant differences in DS+EGCG and DS (intersection), and the number of unique significant differences DS+EGCG *vs* EU (right).

A similar pattern was detected in the 4-12 year age group. EGCG treatment did not produce a substantial change in the number of facial differences between euploid and trisomic children. The baseline comparison of DS and euploid cases estimated that 59% of facial measurements were significantly different in this age-range (Fig. 4e), and this percentage was only reduced to 46% when comparing EGCG treated individuals with DS to the euploid sample (Fig. 4f). However, we detected that the number of significant shape differences between treated and untreated children with DS was larger in the 4-12 age group (Fig. 4g) than in the 13-18 age group (Fig. 4c). This result could be suggesting an incipient positive effect of the EGCG treatment that was however not sufficient to rescue the facial phenotype entirely (Fig. 4g, h).

Finally, the analysis of the youngest age group of 0-3 year olds revealed the most significant effect. The EDMA comparison between euploid and DS children showed that 57% of linear distances were significantly different (Fig. 4i). However, only 25% of linear distances remained significant when young children with DS treated with EGCG were compared to the euploid sample (Fig. 4j). This indicated an overall improvement of more than 50% of total facial measurements relative to the baseline comparison. A significantly large beneficial EGCG treatment effect is thus suggested, especially on those facial traits that appear as significantly different when comparing EGCG treated and untreated children with DS (Fig. 4k, l). The local EDMA morphological assessments indicated that various dimensions of the nose, philtrum, cheeks and midface were improved after EGCG treatment, with the strongest effect occurring in the age group of 0-3 year olds, but with less of an effect in older age groups.

To verify the results suggesting that EGCG improved facial development in young individuals with DS, summary scores of facial improvement (FISs) were estimated and tested using the random simulation approach. The FIS score quantified whether EGCG treatment decreased or increased the number of shape differences identified by the EDMA between children with DS and euploid children. Results showed that in older children (from 13 to 18 and from 4 to 12 years old), the FIS score was positive but low, indicating only a minor relative improvement of facial shape (Fig. 5). The simulation tests indicated, however, that the effect of EGCG at these later stages of development was not significantly different from what would be obtained from a random distribution (Fig. 5). In contrast, the results at the earliest stage of development showed a large positive effect. In 0 to 3 year old children with DS treated with EGCG, the facial improvement was high (FIS=58%) and significant. The simulation tests showed a clear tendency towards increasing the FIS as the number of DS treated cases included in the simulations increased (Fig. 5). Since there were only three random simulations that produced higher FIS scores, our results demonstrate that the effect of EGCG treatment on facial shape at young age is highly significant (*P-value*=0.008) and beneficial.

**Figure 5.**
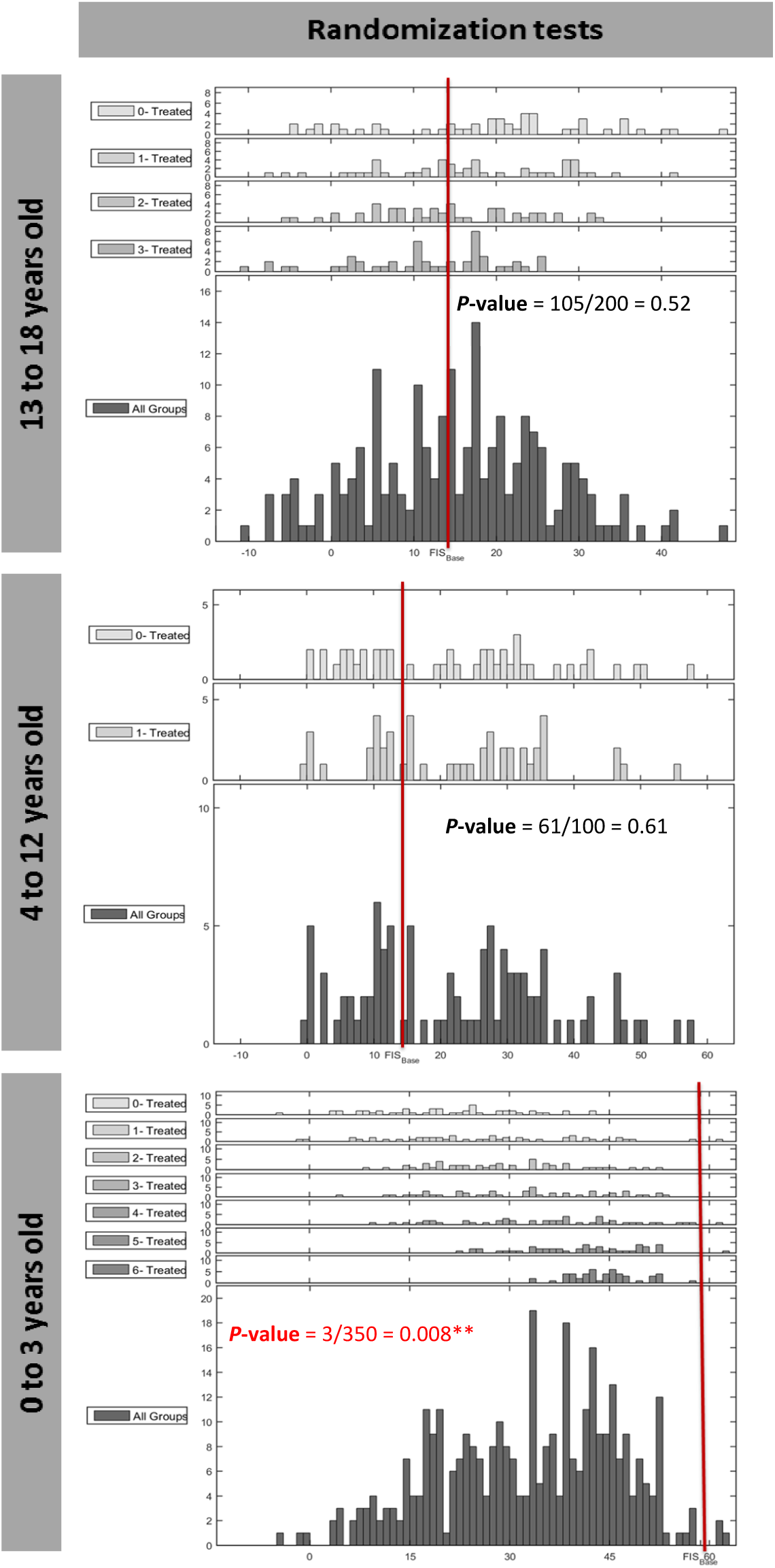
Simulation tests based on facial improvement scores (FIS) at each age group. FIS is computed by contrasting the number of significant differences in facial traits between 1) euploid children and children with DS not treated with EGCG and 2) euploid children and children with DS that have been treated with EGCG. Histograms representing the simulation results for each random group are provided separately as well as aggregately. Each group contains an increasing number of EGCG treated DS cases, from 0 to 6. The red line shows the FIS score obtained with the observed including 7 EGCG treated DS cases. *P-values* are provided for each group, statistical significant comparisons are marked with **.

## Discussion

No approved intervention exists for the clinical amelioration of individuals with DS. However, accumulating evidence is revealing that modulation of a complex regulatory circuit controlling neural, facial and skeletal development by EGCG kinase activity inhibition can improve several aspects of the DS phenotype^21,24^. Previous research has suggested that EGCG enhances cognitive function in individuals with DS^13,16^, as well as cranial morphology^17^ and long-bone skeletal architecture^14,15^ in DS mouse models. Our study extends the evidence for potential benefits of the EGCG treatment to facial morphology with new experimental data on dosage effects, and provides, for the first time, clinical evidence of EGCG effects in an observational study.

## Effects of EGCG treatment depend on the dose in mouse experimental data

Our analyses using *Ts65Dn* mouse model (Fig. 1) suggested that administration of EGCG at low doses from E9 to P29 improved facial dysmorphology. However, the same treatment at high dose produced disparate facial morphology changes with an extremely wide and abnormal range of variation (Figs. 1 and 2). Since some mice receiving the high EGCG dose had relatively normal facial morphology, while others exhibited different and unique dysmorphology (Fig. 1), the bimodal distribution of these results may be suggestive of a threshold effect of EGCG that alters developmental canalization and reaction norms by shifting craniofacial development to a different developmental pathway with a separate phenotypic fate^27^. The unpredictable facial morphologies resulting from high EGCG doses, affecting both wild type and trisomic mice, require further investigation to confirm the potential therapeutic window of EGCG to prevent and alleviate facial dysmorphologies associated with DS.

Ours results agree with previous research using DS mouse model experiments indicating that the potential rescuing effect of EGCG treatment depends on several factors, such as dosage, onset and duration of the treatment, as well as method of delivery and composition of EGCG-containing supplements^19^. De la Torre et al. (2014) showed that 1 month of 2-3mg of EGCG per day can rescue cognitive deficits in 3-month old adult *Ts65Dn* mice (trisomic background including *Dyrk1a* triplication), and transgenic *TgDyrk1a* mice (disomic background including *Dyrk1a* triplication). This result concurred with the effect observed in humans, where the combined intervention of EGCG treatment and cognitive training seemed to ameliorate memory and executive deficits in young adults with DS after taking oral doses of 9 mg/kg/day of EGCG for 3-12 months^13,16^. However, *Ts65Dn* mice treated with ∼10, ∼20, or∼50mg/kg/day of EGCG starting at P24 for 3-or 7-weeks failed to show cognitive improvements^28^.

Regarding the effect of EGCG on skeletal traits, a recent study indicated that EGCG treatment of 200 mg/kg/day on gestational days 7 and 8 of pregnant *Ts65Dn* DS mouse models rescued some cranial vault morphologies measured two months later in 6-week old offspring^17^. Research on limb long bones has shown that specific conditions of EGCG treatment can lead to correction of skeletal deficits associated with DS. For instance, when 3 week old *Ts65Dn* mice were treated with ∼9 mg/kg/day of EGCG for 3 weeks beginning in early adolescence, they exhibited increased femoral bone mineral density relative to untreated trisomic mice and trabecular microarchitecture (*i.e.*, percent trabecular bone volume, trabecular number and thickness) was rescued to euploid levels^15^. However, *Ts65Dn* mice treated with ∼50mg/kg/day for 7 weeks starting at P24 exhibited detrimental effects on femoral bone structure and mechanical properties, such as reductions in cortical area and thickness, structural strength (yield force), stiffness, and ultimate force^19^. Further experimentation with mouse models is thus critical to explain the discordant effects of different conditions of EGCG treatment.

### Potential of EGCG for facial bone remodeling

In the observational study in humans, our morphometric assessment of facial shape revealed the strongest and most beneficial effect for EGCG in the youngest human age group, 0-3 years, when most craniofacial growth occurs (Fig. 4). Only younger children with DS treated with EGCG since their first months of life up to 3 years presented with facial phenotypes that fell close to or within the euploid range of variation (Fig. 3c). It is during these first years of life when the skull and the face exhibit high growth velocities associated with relatively large size and shape change^29^, likely driven by rapid brain expansion and masticatory changes as offspring are weaned to semi-solid and solid foods. After facial shape maturation is reached by 15-16 years of age^29^, facial shape is maintained fairly constant and bone remodeling only occurs at a lower rate. This period coincides with our oldest age group (13 to 18 years), in which no effect of the EGCG treatment was observed, probably because faces have already attained their final size and shape (Figs. 3a, 4a-d) and facial plasticity is minimal.

Since soft-tissues of the face are built upon the bony scaffold of the skull^30^, and because we found soft-tissue morphology improvement in humans and facial skeleton improvement in mouse models using a low dose of EGCG, we suspect that EGCG has a potential bone-remodeling effect during early development and subsequent growth. It has been previously shown that EGCG supplementation can normalize many skeletal deficits. EGCG is known to modulate bone metabolism^31^, bone formation^32,33^, and bone resorption^34^, potentially by hindering osteoclastogenesis^34^ and inducing osteoblast differentiation^35–39^. However, as we found in our experimental analyses on the effect of EGCG on facial shape (Figs. 1 and 2), the specific effects greatly depend on the dosage: high doses of EGCG may suppress bone formation and exacerbate skeletal defects, whereas moderate doses of EGCG may enhance bone formation and attenuate birth defects^40,41^.

We hypothesize that the distinctive faces associated with DS may result from poor craniofacial growth. As individuals with DS show overall impairment of growth and bone formation, displaying shortened limbs, low bone mineral content and density, reduced bone strength^42^ and remodeling^43^, as well as osteopenia and osteoporosis that worsens with age^44^, deficiencies in remodeling and relocation of craniofacial bones as the face grows during the first years of development^45^ may explain the typical facial traits associated with DS. Increased bone resorption, reduced bone deposition, and/or lack of proper relocation may cause midfacial hypoplasia and the altered shape and position of the maxilla, the mandible, and nasal and orbital cavities, which are the bones with higher predisposition to resorption^45^ and are indeed the most affected by Ts21.

Despite all the caveats of an observational study based on a limited sample size, including children from different ages, sexes, and geographic regions and the fact that they did not received the same doses of EGCG or had the same length of treatment, the simulation tests consistently detected a signal indicating that EGCG treatment since early in development is associated with intermediate facial phenotypes and significant facial improvement scores (FISs) (Fig. 5). The findings of our observational study underline the need to reassess the effects of EGCG on facial morphology in a clinical trial, to statistically assess and control the different sources of morphological variation in larger samples of individuals following a systematic treatment.

## Modulating a multisystem circuit to minimize developmental alterations in DS

The pleiotropic effects of EGCG are likely mediated by a complex regulatory circuit controlling neural, facial, cardiac and skeletal development. EGCG likely inhibits the activity of several kinases across different organism tissues^19,46^, including DYRK1A, a protein that is encoded by one of the genes that has emerged to play a critical role in DS. *DYRK1A* is expressed in a gene-dosage dependent manner within a critical region of chromosome 21, and both under-or overexpression of *DYRK1A* lead to neural dysfunction^21^ as well as to brain, craniofacial and skeletal dysmorphologies^17^. The available evidence suggests that *DYRK1A* participates in a regulatory circuit of neural, skeletal, as well as cardiac development, probably mediated through varying control of the Nuclear factor of activated T-cells (NFAT) signaling pathway, a critical regulator of vertebrate development and organogenesis^24^.

The whole circuit is clearly altered in DS by the 1.5-fold increase in gene dosage due to Ts21. Although all the genes and signaling pathways participating in this circuit are yet to be discovered, research on *DYRK1A* has led to the discovery that a multitude of DS phenotypes can be modulated through inhibition of kinase activity by EGCG. However, the potential rescuing effect of EGCG treatment on multiple systems depends on several factors, such as dosage, onset and duration of the treatment, as well as method of delivery and composition of EGCG-containing supplements. Hence, additional research is needed to pinpoint the optimal dose and time window for EGCG administration, minimizing possible detrimental and side effects, and maximizing benefits to cognitive function and biological growth, including skeletal and facial development. This research is critical since EGCG is already commercially accessible and a growing number of people are consuming EGCG as a dietary supplement with no medical prescription or governmental regulation.

## Conclusion

Here we demonstrate the potential of EGCG to alleviate facial dysmorphology in DS, likely through a bone-remodeling effect. The results presented here strongly suggest that supplementation with green tea extracts that contain EGCG during early growth and development improves facial dysmorphology of human children with DS. The experimental studies of *Ts65Dn* mice confirm these effects at low EGCG dose, which corresponds to the one used in individuals with DS^13,16^. However, the appearance of potentially detrimental effects in mouse models when high dose of EGCG treatment are used, emphasizes the importance of identifying dosages of EGCG that reliably improve DS phenotypes as a whole, linking the effects to actions of EGCG in specific targets of the brain, the face and craniofacial bones. Current evidence warns against the generalized, non-prescribed intake of this supplement as a health-promoting measure^47^, as non-tested doses of EGCG can have detrimental effects on both trisomic and euploid conditions.

## Methods

### Mouse experiments

We performed two carefully controlled experiments using the *Ts65Dn* mouse model and assessed the effect of two doses of EGCG treatment on the facial shape of adult mice.

#### Animals

For each experiment we established a breeding colony composed of 4 WT males and 6 *Ts65Dn* females (refs. 001875 and 001924, the Jackson Laboratory Bar Harbor, ME, USA). Mice were housed at the animal facility of KU Leuven in standard individually ventilated cages (40 cm long × 25 cm wide × 20 cm high) under a 12h light/dark schedule in controlled environmental conditions of humidity (50% - 70%) and temperature (22 ± 2°C) with food and water supplied *ad libitum*. Experimental protocols were in compliance with all local, national and European regulations and authorized by the Animal Ethics Committee of the KU Leuven (ECD approval number P004/2016).

#### Treatment

In the first experiment, mice were treated with a high dose of EGCG (30 mg/kg/day), and in the second experiment they received a lower dose (9 mg/kg/day). Since EGCG crosses the placenta and reaches the embryo^48^, the treatment was administered prenatally at embryonic day 9 (E9) to optimize treatment efficiency. We chose this time point because this is when the face starts developing and *Dyrk1A* begins to be expressed in mouse embryos. EGCG treatment continued over the duration of the experiment, from E9 until adulthood at P29. During gestation and lactation, EGCG was delivered to dams dissolved in water at a concentration of 0.326 mg/L in the high dose experiment and 0.09 mg/L in the low dose experiment. After weaning at postnatal day 21 and until postnatal day 29, EGCG dissolved in water at the same concentrations was made available to young mice that drank *ad libitum*. The EGCG solution was prepared freshly from a green tea extract [Mega Green Tea Extract, Life Extension, USA; EGCG content of 326.25 mg per capsule] every 3 days. The dosage received was thus approximately 30 mg/kg/day in the high dose experiment, and 9 mg/kg/day in the low dose experiment for an adult mouse, taking into account that, on average, mice weigh 20 g and drink 2 ml of water per day.

#### Morphometric craniofacial study

We used a total of 55 mice from eight litters. We administered EGCG prenatal treatment to 50% of pregnant dams, chosen randomly, to produce approximately the same number of EGCG treated and non-treated mice. Mice were genotyped by quantitative PCR from ear snip samples and were distributed in four groups according to their genotype and treatment: Euploid (WT), Euploid treated with EGCG (WT+EGCG), trisomic *Ts65Dn* (TS) and trisomic *Ts65Dn* treated with EGCG (TS+EGCG). See Table SI_1 for details on sample composition.

We acquired whole-body micro computed tomography scans of the mice at postnatal day 29 using a dedicated fast low-dose *in vivo* μCT scanner (SkyScan 1278, Bruker Micro-CT, Kontich, Belgium) at high resolution (50 μm isotropic). Free-breathing mice were scanned under isoflurane anesthesia (∼1-2% in oxygen), while physiological functions were constantly monitored and maintained via a feedback loop (temperature, breathing, visual inspection via camera). Scans were reconstructed using the software NRecon v1.6.10.4, whereas skull visualization and landmarking were performed by a single observer using the Amira 6.3 software (Visualization Sciences Group, FEI). Measurement error was within the range of previous studies^49^. We recorded the three-dimensional coordinates of a set of 12 facial anatomical landmarks (Fig. SI_1 and Table SI_2), to capture the shape differences between WT and TS mice and to assess whether the EGCG treatment has an effect on facial development.

### Patient cohort and recruitment

#### Facial image acquisition and anatomical landmark collection

The human sample includes 288 European and North American children from 0 to 18 years old recruited for photographic sessions at schools, research centers, and DS family congresses between 2009 and 2017 (Table SI_3 and SI_4). To photograph the children and to obtain relevant clinical information we obtained informed consent from their parents or legal tutors (CEIC-Parc de Salut MAR, N° 2012/4849/I; UCF IRB approval SBE-15-11524). Human faces were digitalized using non-invasive photogrammetry techniques. The facial images of each individual were acquired in an upright position with neutral facial expression using either a 3dMD Face system (3dMD, Atlanta, GA) or self-built photographic rigs, as previously described (Starbuck et al. 2017, González et al 2018). Each system acquired 5-10 images simultaneously of an individual’s face and used photogrammetry to merge separate two-dimensional images into a single three-dimensional model that accurately represents facial shape. The software used for obtaining the 3D reconstructions was 3dMD (3dMD, Atlanta, GA), PhotoModeler (2015) version 1.1.1546 (Eos Systems, Inc. Canada), and AgiSoft PhotoScan Standard (2016) version 1.3.2. All data was collected with high precision and accuracy based in the same principles of stereo photogrammetry, thereby avoiding potential bias due to the use of different photographic set-ups and software. Scale differences from the different acquisition systems were accounted for in the subsequent analyses by transforming pixel sizes to their corresponding mm measures.

We collected 3D coordinates of 21 facial landmarks from each individual’s 3D image following published protocols^7^. Landmarks correspond to homologous biologically meaningful points with precise anatomical definitions that can be reliably registered across all individuals (Fig. SI_1 and Table SI_5). Measurement error was within the range of previous studies (0.26 mm along the x-dimension, 0.30 mm along the y-dimension, and 0.31 mm along the z-dimension)^50^ and thus considered negligible. Following inspection of landmark coordinates for gross error, anatomical landmark coordinates were used to conduct morphometric analyses.

### Morphometric analyses

We compared facial shape using different geometric morphometric methods.

#### Procrustes method

First, we used a General Procrustes Analysis (GPA)^25^ to assess the facial phenotype from a global perspective, considering facial shape as a whole. Using this method, the 3D spatial relationship between landmarks defining facial shape was always preserved throughout the analysis, and facial shape was analyzed as a whole 3D facial shape landmark configuration. Within the GPA, landmark facial shape configurations of all individuals were superimposed by shifting them to a common position, rotating, and scaling them to a standard size until a best fit of corresponding landmarks was achieved. Although the resulting Procrustes coordinates were scale-free, the shape data still contained a component of size-related shape variation due to allometry^51^, which refers to organism shape change relative to body size changes that occur from growth during development. Since our human dataset included children of different ages, we removed the allometric effects by performing a multivariate regression of shape on age within each age group^51^. The Procrustes residuals from this regression were the input for further statistical analysis.

Principal Component Analysis (PCA) was used to explore facial morphological variation^25^ and to test whether individuals grouped according to their genotype and/or treatment. PCA is a multivariate technique that performs an orthogonal decomposition of the data and transforms variance covariance matrices into a smaller number of uncorrelated variables called Principal Components (PCs), which successively account for the largest amount of variation in the data. Each individual is then scored for every PC, and individuals are plotted using these scores along the morphospace defined by the principal axes as an ordination method to visualize patterns of shape variation associated with individual axes. The distribution of groups throughout the morphospace along different PC axes reveals similarities or differences in facial shape and variation among groups. Importantly, this ordination method allowed us to assess whether individuals with DS treated with EGCG present an intermediate facial phenotype relative to untreated individuals with DS and euploid controls along the PC axes explaining the most variation across groups.

#### EDMA method

EDMA is a powerful morphometric method for assessing local differences between samples because it allows researchers to pinpoint exactly which linear measurements significantly differ between pairwise sample contrasts and to compare patterns of significant differences across samples^26^. In the human sample, using EDMA two-sample shape contrast analyses based on a non-parametric bootstrap (10,000 resamples), a total of 210 unique facial measurements were calculated from Procrustes-based allometrically corrected shape coordinates and evaluated for each age grouping (0-3 yrs., 4-12 yrs., 13-18 yrs.) of the DS, DS treated with EGCG, and euploid groups. Local EDMA results, based on confidence interval testing (α = 0.10), revealed unique morphological patterns of variation for each two-sample comparison by evaluating the number of statistically significant linear distances in each comparison and by plotting significant linear distances upon facial figures to pinpoint specific local shape differences.

#### Facial improvement score (FIS)

Using the results of the EDMA analyses we estimated a relative facial improvement as the following percentage:

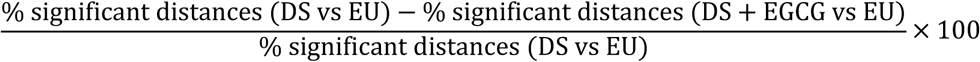

To verify the statistical validity of the patterns revealed by the EDMA analyses, we ran simulation tests. We performed a subsampling approach in which we generated random pseudo-subsamples with increasing number of DS treated cases. For each age group, the whole sample was randomly divided into groups of N individuals (where N is the number of EGCG treated individuals with DS in the original sample) following a controlled random subsampling approach that created pseudo sub-samples containing a known number (namely M) of treated individuals. Thus, a series of random pseudo sub-samples containing from M=0 to M=N treated individuals were created. We computed a FIS score corresponding to each simulation and analyzed the FIS score histogram of each random group. Finally, we computed an aggregated FIS score histogram of all groups and compared the distribution of random FIS scores with the observed FIS score. The *P-value* of the comparison was computed as the ratio between the number of simulations that provided a higher FIS score than the observed FIS score and the total number of simulations and was used to assess statistical significance (α = 0.05).

## Acknowledgements

We gratefully acknowledge the participation of all the children and their families, as well as all the Down syndrome family associations supporting this study. We are indebted to Max Rubert for his continuous technical photographic assistance. This was a project supported by a 2016 Leonardo Grant for Researchers and Cultural Creators, BBVA Foundation, to NMA. The Foundation takes no responsibility for the opinions, statements and visual content of this video, which are entirely the responsibility of its authors. We acknowledge further grant support from the following funding agencies: CRG Awards for Collaborative Research Proposals 2015 to NMA and JA, American Association of Physical Anthropologists Professional Development Program Grant to JMS, Flemish Research Foundation (FWO) fellowship KU Leuven IF STG/15/024 to GVV, and MINECO SAF2016-79956-R to MD. We acknowledge support of the Spanish Ministry of Economy and Competitiveness, ‘Centro de Excelencia Severo Ochoa 2013-2017’, SEV-2012-0208.

## Author contributions

JMS, MD, GVV and NM-A conceived the project. JS, MD, GVV and NM-A contributed with experimental mouse models; SL, JA and JW conducted experiments and scanning; JMS, RdT, MD and NM-A recruited participants for observational study; JMS, SL, RG, AS, AG, XS and NM-A collected and analyzed the data. NM-A and JMS wrote the paper and all authors critically reviewed the manuscript.

